# Reconstructing the decline of Atlantic Cod with the help of environmental variability in the Scotian Shelf of Canada

**DOI:** 10.1101/2022.05.05.490753

**Authors:** Jae S. Choi

## Abstract

Ignoring environmental variability can lead to imprecise and inaccurate estimates of abundance and their spatial distribution of organisms. Fully embracing environmental variability can improve precision and accuracy of estimates of abundance and distribution, especially when they can often be measured with lower costs. Using the example of Atlantic cod in the Scotian Shelf of the northwest Atlantic Ocean, we demonstrate the improved clarity of their historical population trends when such informative features are included. Further, the use of Bayesian spatiotemporal Conditional auto-regressive models substantially improves our ability to understand the role of ecosystem variability upon cod, even when samples are incomplete or missing. Finally, by decomposing biomass into number, weight and a Hurdle process to estimate habitat conditions, we can extract much more information on what has occurred in the past and make reasoned inference on processes.

**One-Sentence Summary:** Deconstructing and reconstructing cod with environmental variability

## Introduction

Environmental variability is almost always perceived as a nuisance or obstacle that hinders the precise and accurate estimation of the abundance of organisms. The perspective originates from the historically dominant influence of Analysis of (Co)Variance approaches where one attempts to “block out” such uncontrolled or uncontrollable factors, using some form of a stratified random design. This is, of course, a powerful approach that has supported much of scientific discovery. But key to the successful application of such an approach is that the stratification must correctly block out the factor(s) that interfere/influence such that samples from within a given stratum (and usually, between stratra) are independent and identically distributed (IID). In a large marine spatiotemporal context, this requirement generally can not be guaranteed due to the multiplicity of space-time scales in operation. Examples of important ecosystem features known to influence biota include: oceanic currents, tidal currents, global climate variability, temperature variability, salinity variations, light levels, nutrient/resource availability, predator-prey interactions, disease, etc. Defining strata that will ensure such features are homogenous within an areal and temporal unit is remarkable difficult to accomplish. As environmental variability is spatially and temporally complex and do not respect the fixed area and time definitions that we might impose, adoption of a naive and large stratified random design can create estimation problems.

Historically, the Atlantic cod in our focal area of study, the Scotian Shelf in the northwest Atlantic Ocean, was sampled under such a stratified random design with fixed areal bounds and sampled loosely as “Summer” and “Winter” time periods [1]. The areal units were fixed at the onset of the survey series in 1970 and utilized depth and oceanic current information available at the time to delimit the bounds (Fig 1). The assumption was that samples within each, generally very large areal (and temporal) unit, would be IID and that between areal (and temporal) unit variability could be treated as independent factors. At the time of inception of the surveys, information on the system was limited. Depth was used presumably because it is a well-known proxy for pressure, light, turbulence, substrate complexity, and overall variability of any number of other environmental factors (temperature, conductivity, salinity, pH, etc.) and so permits some measure of implicit “control” over these factors.

**Fig 1.**
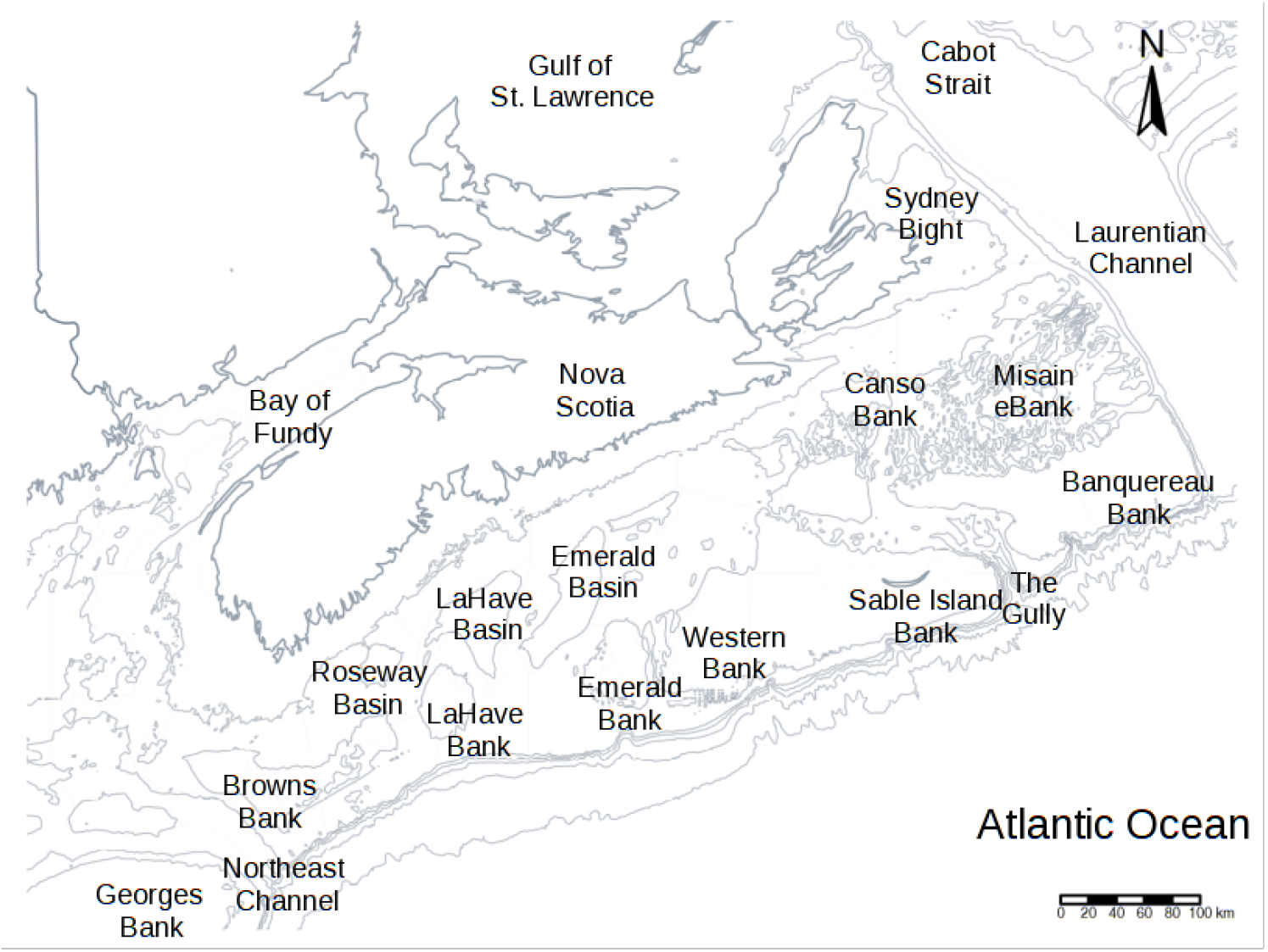
The area of interest in the Scotian Shelf of Canada, in the northwest Atlantic Ocean. Shown are isobaths and some of the major bathymetric features in the area. It is hydrodynamically complex due to the confluence of cold, low salinity water from the north with warm saline water from the south, frequently disrupted by multi-scaled climatic forcings.

Unfortunately, the numerous relevant ecosystem features are not static nor does depth fully control them. This is particularly the case in the focal area as it is at the confluence of the Nova Scotia Current that flows southwest along the shoreline, the Gulf stream that flows northwest along the shelf edge, the Labrador current that flows southwest along the shelf edge and coast waters, and the Gulf of St-Lawrence low salinity waters that flow southeast; it is hydro-dynamically very complex, influenced by global- and regional-scaled climatic variability [2]. Naive random stratification essentially amounts to hoping the fixed area-based design will block out ecosystem variability, both within areal and temporal units. These factors being dynamic and spatially structured, are not homogenous within nor between areal units. Instead, they are entangled across multiple scales of space and time and so complexly correlated.

Here we suggest that, instead of seeing such biologically important ecosystem features as a nuisance or problem to be abstracted away, if we instead embrace them, they can actually increase the precision and accuracy of estimation. This is because, many such ecosystem factors can now be measured at far greater data density than the primary biological samples. The former generally have relatively lower costs, or preexist in multiple monitoring programs or accessible through remote sensing technology. Indeed, they can help fill in the gaps of sampling and even condition/constrain interpolations or extrapolations (such as when surveys do not complete due to operational breakdowns).

Further, a given areal unit has some similarity in environmental and biological processes to a neighboring areal and temporal unit, a similarity that generally declines with distance or connectivity and time. As stated by Tobler’s First law of geography [3]: everything is related to everything else, but near things are more related than distant things. If it were not the case, then everywhere we look would be completely chaotic and discontinuous, without localized areas of homogeneity. Similar arguments can be made for time. Rather than ignoring this process by treating areal units and time as two independent factors, we can also embrace spatiotemporal autocorrelation. Here, we focus upon a simple method that is a minor extension of Generalized Linear Models known as Conditionally AutoRegressive (CAR) models because they have been shown to be both robust and effective even when sparse spatial distributions exist associated with large declines in abundance, yet versatile enough to permit modeling of important covariates in a fully Bayesian context.

A direct analysis of biomass using CAR is possible, however, much more information can be extracted that relates to it by decomposing it as the product of weight and number. This decomposition is especially useful as population dynamic models generally focus upon numbers rather than biomass. Further, the mean number can be decomposed through the Hurdle process [4,5] into a binary component, below or above some detection limit and so characterizes the probability of observing or not observing an individual beyond some threshold; and a numerical count once this “hurdle” is passed. The binomial component was examined elsewhere [6] and how cod habitat quality deteriorated over time. This paper builds upon this and combines the CAR/Hurdle approach to decompose the history of Atlantic cod and then reconstruct the biomass dynamics over the historical record with the goal of understanding this persistent decline.

## Methods

For these analyses, we drew upon two high-quality bottom trawl-based research surveys conducted in the Northwest Atlantic Ocean of Canada. The first is an annual multispecies survey conducted by the Department of Fisheries and Oceans (DFO) from 1970 to the present (“ground fish survey”). Sampling stations were randomly allocated in depth-related strata with sample tows of approximately 1.82 to 2.73 km in length on a variety of different gears, mostly the Western IIA net. Vessel speeds ranged from 4.5 to 6.5 km/hr. Only tow-distances were available which was used to approximate areal density using a nominal wing spread of 12.5 m. Other sampling gear have been used over the years. From 1970 to 1981, the Yankee 36ft trawl was used, which for cod is assumed to have an efficiency of 80% relative to Western IIA. This crude “correction factor” was not used in this analysis; the naive depth-stratified random sampling design (“Stratanal”; Fig 2, blue line) from 1970 to 1981 are thus, marginally lower than what would have been obtained if the correction factor had been used.

**Fig 2.**
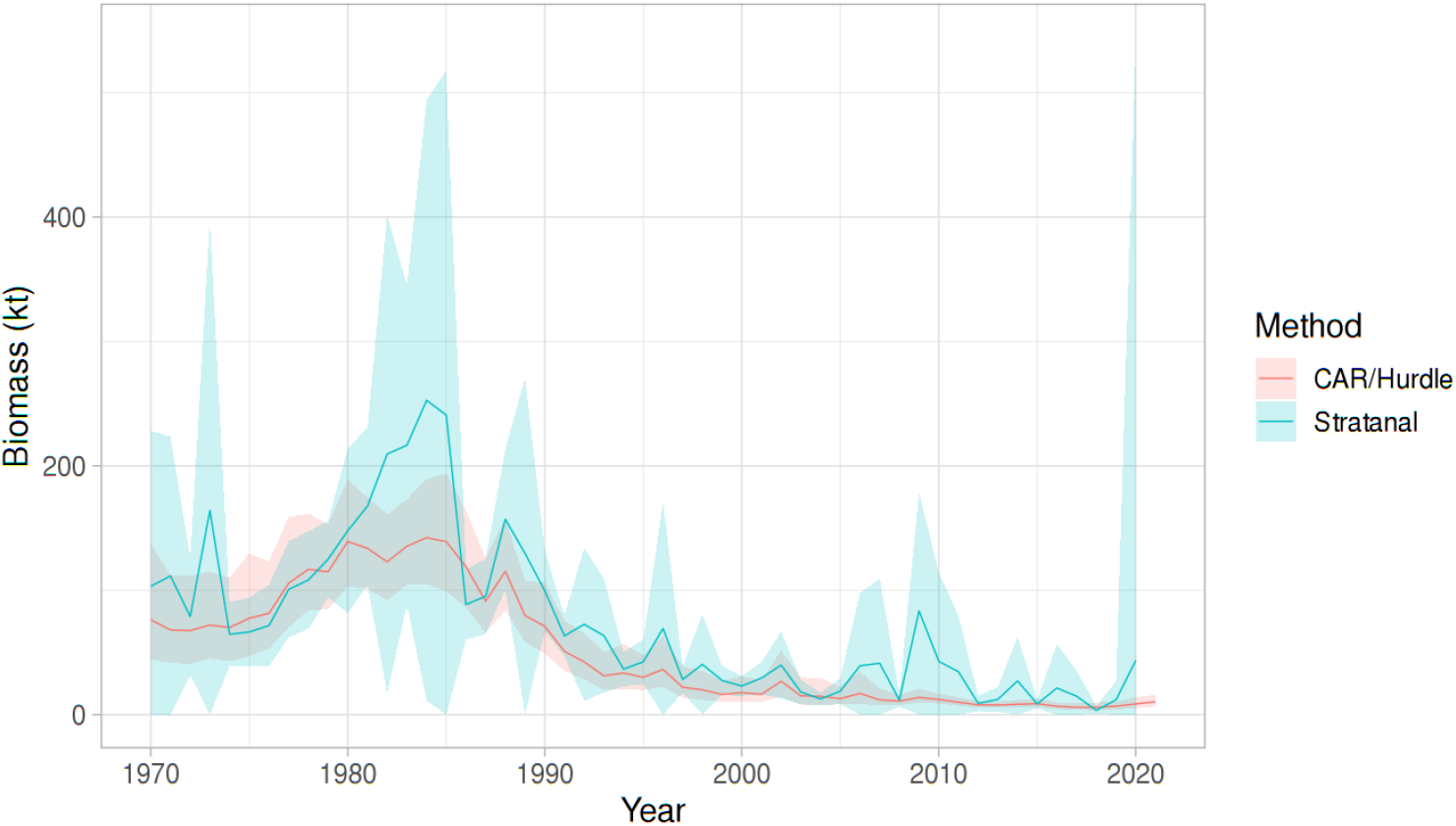
Biomass of Atlantic cod (kt) in the “Summer strata”. The 95% Confidence Intervals from the Stratanal approach to biomass estimation (blue; asymptotic t-distribution assumption) and the posterior predictors for the CAR/Hurdle method (red). Note the large inter-annual fluctuations and large Confidence Intervals that often cross the zero-line. In contrast, note the much smaller Credible Intervals for the CAR/Hurdle estimate. Ground fish surveys were incomplete in 2018 and 2021; associated Stratanal estimates are not comparable. From 1970 to 1981, a Yankee 36ft net was used that is assumed to have 80% net efficiency relative to the net used from 1983 to 2022. This correction factor is not used in the Stratanal estimate.

The second data series is a survey conducted through a joint collaboration between the snow crab industry in the region and DFO, from 1999 to the present (“snow crab survey”). The survey is funded by a proactive industry as a forward-looking investment towards long-term sustainable fishing, an exemplary model of the precautionary approach to fishing. A modified Bigouden Nephrops net designed to dig into sediments was used with sample tows of approximately 0.3 km in length and a target vessel speed of 3.7 km/h. Bottom contact and net configuration were monitored to provide explicit areal density estimates. The wing spreads ranged from 12 to 14 m, depending upon the substrate encountered.

The procedure to estimate the biomass of Atlantic cod was as follows. The average body weight of cod (M_st_) at areal unit s=(1,…, a) and time unit t=(1,…, b), was modeled as a Gaussian process, where a sampling event had 30 or more individuals to provide reasonably stable estimates of mean weight m_st_, and standard deviation, s:

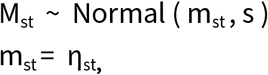

that is, an identity link for the linear predictors η_st_. The linear predictors are defined below.

A binomial model of presence-absence (Y_st_) was developed using the 0.05 probability of the empirical quantile of numerical density (density_0.05_) as a threshold for categorization as presence-absence for a given sampling gear (or alternatively, as a detection limit of the sampling gear censoring observations; see [6] for more details). The binary categorization of each sampling event, Y_st_, is modeled as a *Bernoulli* binomial process with a *logit* link function:

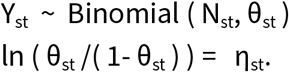

Here, θ_st_ is a vector of probabilities of success (density > density_0.05_) in each trial in each space and time unit and N_st_ the number of trials.

Subsequently, the number of cod counted (D_st_), passing the “hurdle” of density_0.05_, were modeled as a Poisson process:

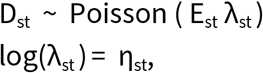

where E_st_ is an offset (swept area and sub-sampling factor) and *λ*_s_ is the rate.

Biomass was estimated as the product of the posterior simulations (n=5000) of mean number and mean weight for each areal and time unit and then corrected for the censoring (truncation; [5]). This separation of the binomial process from a counting process (Hurdle process) is useful when the Poisson process by itself does not capture all the zero-valued catches properly and/or when there is a detection limit or censoring. Negative binomial processes could be used as well if variance was over-dispersed, but here, we limit the analysis to the Poisson case as it captures the variability well.

Each of the above components were modeled, following [6,7]:

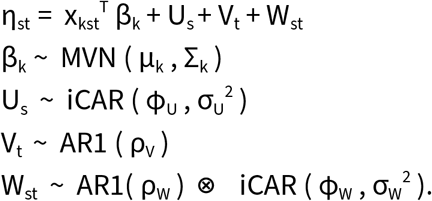

The x_kst_ represent the 3D matrix of covariates with parameters (coefficients) β_k_ distributed as a multivariate normal with mean μ_k_ and diagonal variance matrix Σ_k_; the spatial autocorrelation U_s_ component follows the iCAR [6–8] with ϕ_U_ the proportion of the iCAR variance, σ_U_^2^, attributable to spatial processes relative to nonspatial processes; the temporal error components V_t_ are assumed to follow a lag-1 autocorrelation process (AR1) with autocorrelation parameter ρ_V_; and finally, the spatiotemporal errors W_st_ modeled as a Kronecker product (⊗) of temporal AR1 and spatial iCAR processes, where time is nested in space. This parametrization is known as an “inseparable” space-time model. Amongst the *k* covariates in this study were: bottom temperatures, depth, gear type, season, the first two axes of a species composition ordination (Principal components analysis) and a global intercept term. The Western IIA was used as the reference gear, the longest running survey gear.

As multiple surveys with different sampling designs were used, a hybrid, Voronoi triangulation and tessellation-based solution was used. The algorithm maintains an approximately constant station density within each areal unit, a target of 30 sampling events within each areal unit and a final total number of areal units of less than 2000 units. This was a trade-off between information density and computational stability. The tesselation-based approach had the advantage of more informative areal units capable of resolving temporal trends and environmental gradients. This was critical as environmental variability has been significant in the area/time.

All analyses were conducted with R [9]and R-INLA [10] with all code available online (https://github.com/jae0/aegis.survey/blob/master/inst/scripts/{10a_cod_groundfish_strata. R, 10b_cod_carstm_tesselation.R}). Note that we utilize the “bym2” parametrization of the iCAR for stability and interpretability of parameter estimates [8]. Penalized Complexity (PC) priors that shrink towards a null (uninformative base model) were used for all random effects [11]. For the AR1 process, we used the “cor0” PC prior which has a base model of 0 (i.e., no correlation). For the spatial process (“bym2”) we use the PC prior (rho=0.5, alpha=0.5).

Seasonality was discretized into 10 units and a cyclic “rw2” process was used. For all other covariates, a second order random walk process (“rw2”) was used as a smoother using 11 quantiles as discretization points.

## Results and discussion

The naive depth-stratified random sampling design based time trends of biomass (“Stratanal”; Fig 2) has been the basis of almost every research program related to demersal and benthic organisms in the region since the 1970s. They are also problematic. Being simple area-weighted expansions of the mean biomass, adding other survey data or other gear types requires explicit experiments and modeling. Further another important limitation is that when an areal unit is not visited or there are missing years due to operational issues (weather, vessel break down), there is no recourse. There is no estimate for that areal unit or year and so, they can only be assumed to be zero-valued for a non-surveyed area or some other value in an ad-hoc manner. This has implications for accuracy and precision. If an appropriate sampling distribution is assumed (t-distribution given the small number of samples within each areal unit), poor precision is evident. Further, as each areal unit is assumed to be IID, that is uncorrelated, counter to Tobler’s law. Finally, environmental variability is completely ignored. The temperature and species assemblages (predators, prey, competitors) in the environment in 2020s are known to be very different from those in the 1970s [12], yet as far as the Stratanal approach is concerned, there has been no change. It was, however, an operational approach considered functional at the time of inception, especially when no additional information or resources were available to improve upon the precision and accuracy.

Importantly, there is a lot more additional information on ecosystem variability that has accumulated since the 1970s. We can draw upon them as a source of additional constraints and information through a model-based approach. Indeed, the CAR/Hurdle model approach helps to improve overall precision and accuracy (Fig 2). The immediate outcome are reduced inter-annual fluctuations (and biologically more reasonable), and more credible/believable variability estimates. Further, instead of implicitly assuming areal units that were not sampled to be zero-valued, the CAR/Hurdle approach treats them as random effects that can be estimated with the information gain and constraints from adjacent areas, times, covariate effects and marginal distributions. More consistent comparisons between years can be made, and even reasoned inference of aggregate biomass when surveys do not complete (Fig 2; see year 2021) or when alternate surveys or gear types are used (Fig 3).

**Fig 3.**
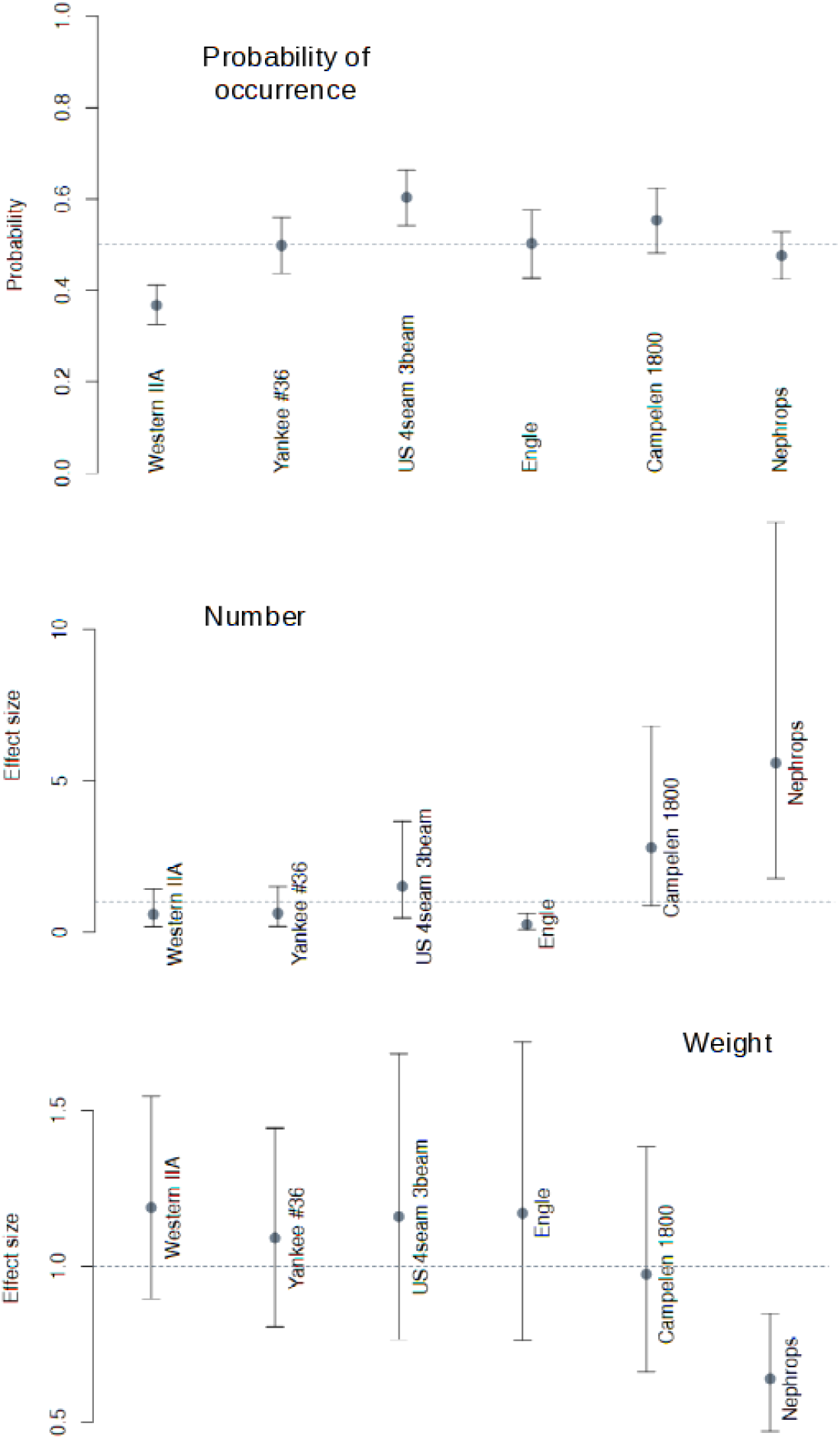
The posterior mean effect size of gear type for each of the models (Hurdle probability, number and mean body weight). The 95% Credible Intervals are also presented.

The CAR/Hurdle analysis suggests biomass levels peaked from 1980 to 1985, a few years prior to that suggested by Stratanal. The magnitude of the peak in biomass may have been much less than previously thought. Indeed, Stratanal seems to significantly overestimate biomass in 1973, 1981-1985 and 2009-2011 (Fig 2). The very high Stratanal estimates would have painted an over optimistic picture of cod abundance in those periods and also their potential intrinsic biological rates (production). Ultimately, over optimistic fishery advice was likely to have been provided. Further, the CAR/Hurdle estimates suggest that the decline was not a dramatic and catastrophic collapse, but rather a gradual decay to an alternate steady state (attractor). Biomass peaked from 1980-1985 and by 1992, the CAR/Hurdle biomass was about the same magnitude as those prior to 1975. After the moratorium, biomass levels continued to decline to a low but steady state.

Decomposing the dynamics of biomass in terms of weight, number and the hurdle (habitat) probability, we see first, a continued degradation in the probability of observing Atlantic cod (i.e., habitat probability) since the mid-1980s (Fig 4). Numbers also peaked a little in 1983 and then declined to less than 40 × 10^6^ individuals by 1993, after which it has been more or less stable. As a result, the decline of biomass in the post-moratorium period is mostly attributable to a decline in average body size from about 1kg in 1992/3 to less than 0.4 kg by 2010, and not number. A much wider range of body sizes were also found in 1978 which has shrunk to extremely low and worrisome level of variability by 2012. The overall decline of almost 300% in average body weight is indicative of strong size truncation and associated population dynamic consequences (lower egg production, susceptibility to replacement by other species due to a lack of dominance). Size at age has also been shown to have declined over these years [13]. Demographically, cod has been exhibiting signs of an alternate attractor due to population level distress and perturbation starting from the early 1980s [13,14].

**Fig 4.**
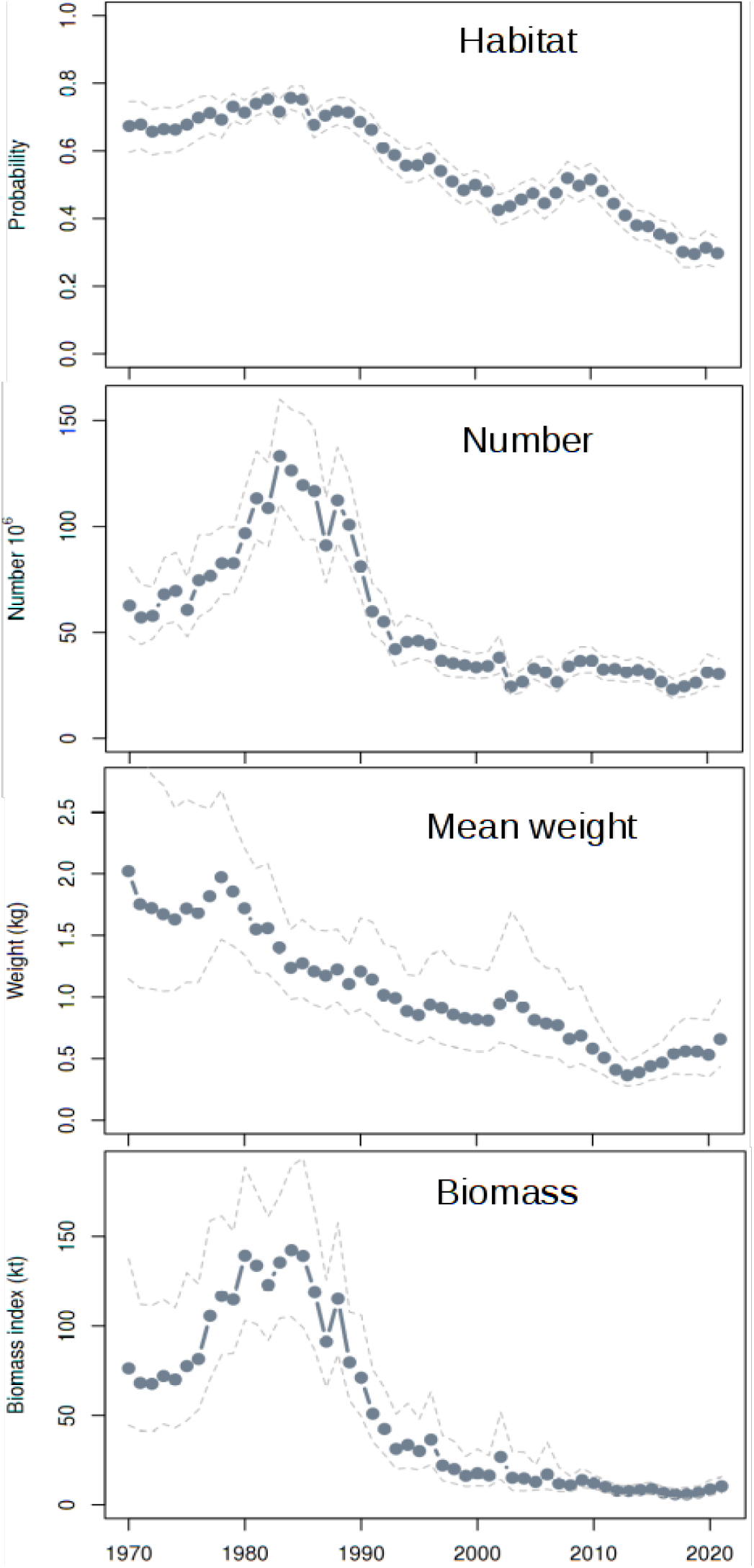
Time series of aggregate indices of Atlantic cod from posterior mean predictions and 95% Credible Intervals. Note: 1981 was a peak year in terms of depth-temperature habitat effects. Years 1989-1992, 2003, 2012, 2018 and 2021 were years with extremely poor depthtemperature related habitat effects (surface area less than 65,000 km^2^) with Credible Intervals non-overlapping with those from peak years (1981, 1984).

Body condition, fishing, predation, and other sources of mortality likely had roles to play in this decline [1,15–17]. However, habitat analysis identified 2012, 2018, and 2021 as having been extremely poor environmental conditions, in terms of bottom temperature-depth effects upon cod habitat [6]. This effect when represented in terms of habitat surface area (where the probability of observing cod due to temperature and depth effects are greater than 0.25; Fig 4) demonstrates the continued shrinkage of habitat in the historical record. We note that 2012 was the year in which mean weight of cod declined to a low of less than 0.4 kg with almost no variability around the posterior mean (Fig 5). Environmental variability also played a role in this decline. The systemic changes noted in the Scotian Shelf suggests that large-scaled processes are involved, and rapid climate change had an important role to play in the decline of Atlantic cod.

**Fig 5.**
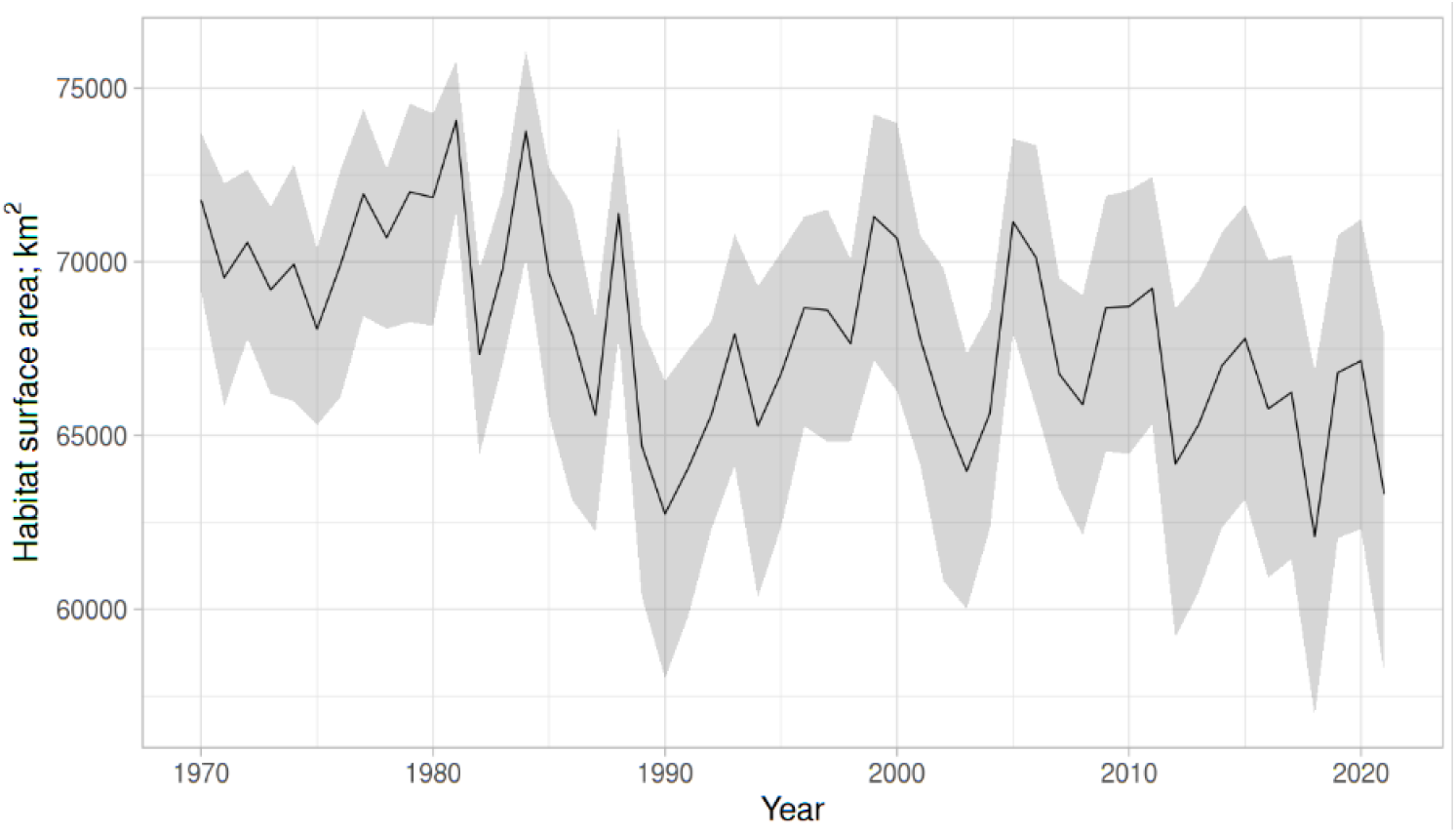
Posterior distribution of Atlantic cod habitat surface area (km^2^) associated with the interplay of depth and bottom temperature spatiotemporal patterns. The area is based the joint probability attributable to depth (for the highest probability subset: 10-100 m) and temperature, being greater than 0.25. Though the overall pattern is not linear, the linear trend is a loss of about 100 km^2^ per year; p<0.0001. Note the declines (below 65000 km^2^) in 1990, 2003, 2012, 2018 and 2021.

Contrasting the spatial variations of cod characteristics between a year with very good habitat conditions (1981) to a year with very poor habitat conditions (2012), we can see the locations where the degradation in body size has been occurring (Fig 6; the year 2021 is also shown for reference) in areas most affected by the high bottom temperatures: Bay of Fundy, Browns Bank and most of the southwest and to the northwest near Banquereau and Western Banks. Numerical and biomass declines were domain-wide. Indeed, at present, only the area near Sydney Bight and Canso Bank have habitats that can be considered reasonably positive for cod (Fig 6). We note that bottom temperature is markedly different between the two years. The bottom temperatures were more moderate in 1981, whereas, in 2012, they were very warm in the southwest from Gulf Stream incursions. The years 2018 and 2021 were even more extreme, but sampling was unfortunately incomplete. The expectation is that in 2022, there will be severe size truncation again.

**Fig 6.**
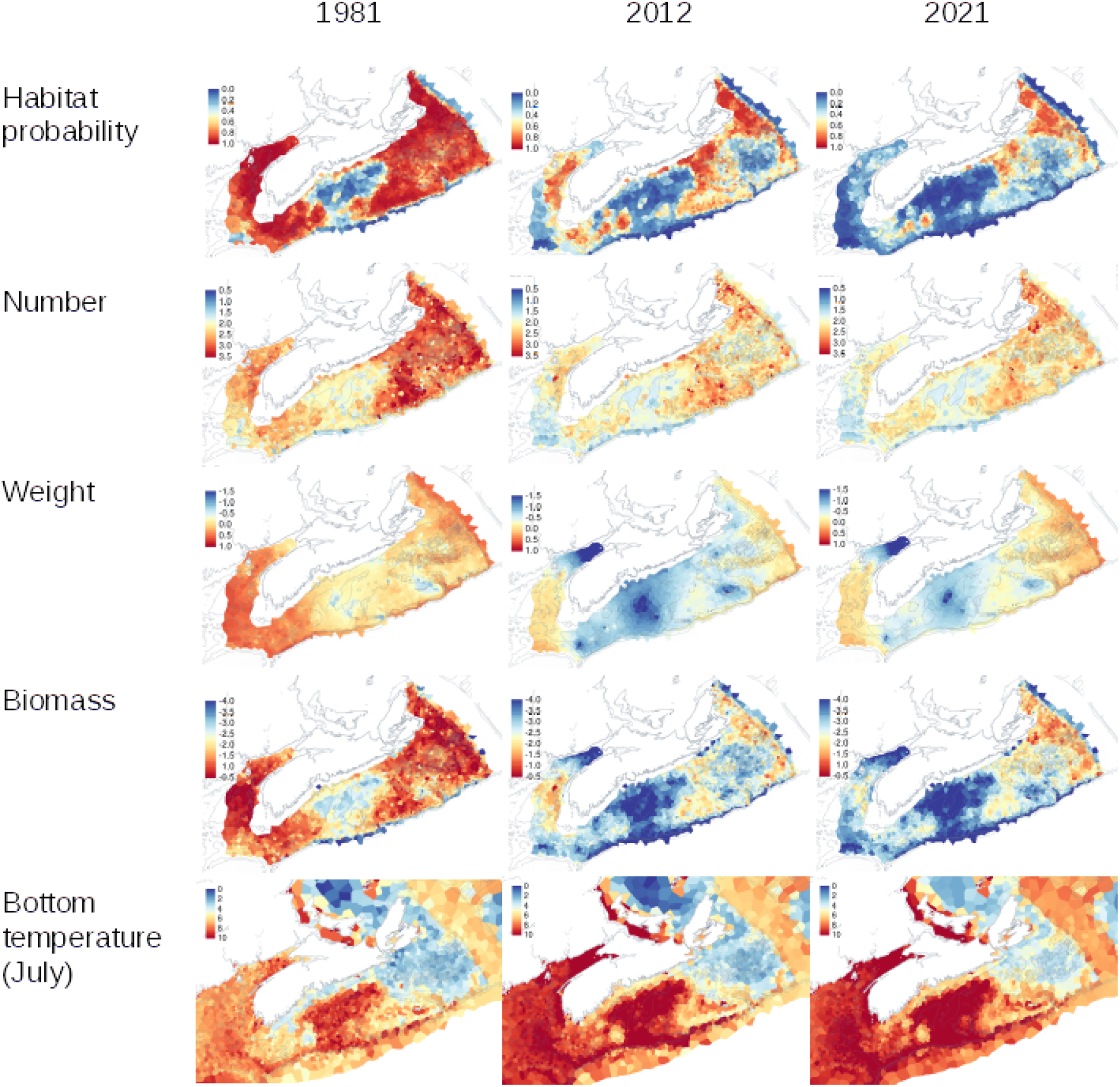
Posterior predictions of the Atlantic cod characteristics and bottom temperatures for two key contrasting years of habitat conditions (1981 vs 2012) and the current state (2021). Note: number, weight and biomass are on log10 scales of areal density (per km^2^) and temperature in degrees Celsius.

## Conclusions

The Car/Hurdle approach is admittedly a simple model-based approach. More complex and powerful approaches exist such as continuous Gaussian Process models (e.g., stochastic partial differential equation approximations). They come a costs of computational time and complexity. The Car/Hurdle approach being simple also facilitates operational use and a reasonable first step in incorporating ecosystem variability into estimation abundance and spatiotemporal distribution of biota.

Environmental variability is important in the Scotian Shelf. It does interfere with estimation of abundance if one tries to ignore it. But using it to inform model constraints and estimation helps to improve precision and accuracy. Decomposition into weight, number and habitat components increases our ability to understand the nature of the changes. Environmental variability cannot be safely ignored especially as it has been an important factor in the decline of Atlantic cod in the area. Future work will continue to add more environmental factors towards understanding the abundance and distributional patterns of Atlantic cod and other species. Already, these methods form the basis of the assessment of snow crab, another important species in the area.

## Data and materials availability

All data are held in databases in Fisheries and Oceans Canada. All analytical code are available at https:://github.com/jae0/aegis.surveys/.

## Reviews

Comments and reviews were kindly provided by Y. Wang, K. Christie, A. Glass, A. Vezina, and M. Greenlaw. All errors are my own.

The perspectives outlined here are those of the author and not necessarily that of funders nor Fisheries and Oceans Canada.

